# From Parametric Guessing to Graph-Grounded Answers: Building Reliable ChatGPT-like tools for Plant Science

**DOI:** 10.64898/2026.04.02.716042

**Authors:** Manoj Itharajula, Shan Chun Lim, Marek Mutwil

**Affiliations:** Interdisciplinary Graduate Programme (AI-X), Graduate College, Nanyang Technological University, Singapore; School of Biological Sciences, Nanyang Technological University, 60 Nanyang Drive, Singapore, 637551, Singapore; Department of Plant & Environmental Sciences, University of Copenhagen, Frederiksberg C 1871, Denmark

## Abstract

Large language models (LLMs) are increasingly used by plant biologists to summarize literature, generate hypotheses, and interpret experimental results. However, LLMs are unreliable sources of exhaustive, source-attributed facts, a critical limitation for the list-style queries that pervade plant biology (e.g., "list all transcription factors regulating secondary cell wall (SCW) biosynthesis in Arabidopsis"). Here, we query ChatGPT, Claude, and Gemini with such queries and demonstrate that none return complete gene lists with reliable citations. We trace these failures to how LLMs store knowledge: as statistical patterns distributed across billions of internal parameters, with no mechanism to guarantee completeness, provenance, or reproducibility. We also review fine-tuning mitigation strategies, including multi-task instruction tuning, parameter-efficient methods, and context engineering, that alleviate but do not resolve these limitations. We then discuss retrieval-augmented generation (RAG), which feeds relevant documents to the LLM at query time, and argue that while it improves source attribution, it remains impractical when answers require synthesizing information scattered across hundreds of papers. As an alternative, we advocate graph retrieval-augmented generation (GraphRAG), in which the LLM serves as a reasoning and language interface over a structured, provenance-linked knowledge graph (KG) that returns complete result sets reproducibly. We outline a practical GraphRAG architecture and survey existing plant KG resources. Finally, we discuss open challenges, including entity disambiguation, relation normalization and evidence grading, and propose a roadmap for building open, continuously updated plant KGs that can turn "read 1,000 papers" into a single reproducible query.

## Introduction

Large language models (LLMs) such as ChatGPT, Claude, and Gemini have captured global attention for their ability to engage in human-like dialogue and synthesize information across a wide range of topics. In life sciences, researchers increasingly turn to LLMs for tasks such as summarizing literature, generating hypotheses, and interpreting experimental results. However, when scientific inquiries demand exhaustive factual enumeration or up-to-date domain knowledge, for instance, "list all transcription factors regulating secondary cell wall (SCW) biosynthesis in Arabidopsis", current LLMs fall short. They offer no guarantee of **completeness** or accuracy: 1) they may omit relevant entities; 2) fabricate plausible-sounding but incorrect facts (a phenomenon termed hallucination); and 3) do so with the same confident tone used for correct answers (Huang *et al*., 2025; Santana Rizzi *et al*., 2025). Crucially, LLMs also lack robust **provenance**; even when citations are provided, studies have shown that 30–90% of LLM-generated references are partially or entirely fabricated (Wu *et al*., 2025a).

These limitations are especially consequential for plant biology. Compared to biomedical science, plant science is underrepresented in LLM training corpora, meaning that model coverage of plant-specific genes, pathways, and regulatory networks is likely to be shallower and less reliable. Moreover, the questions plant biologists routinely ask: 1) which genes belong to a given family across species; 2) which transcription factors regulate a particular pathway; and 3) which protein–protein interactions have been reported, are precisely the list-style, multi-entity queries at which LLMs perform worst.

In this review, we argue that the solution is not to abandon LLMs but to redefine their role: from unreliable knowledge stores to powerful reasoning and language interfaces operating over structured, provenance-linked knowledge graphs (KGs). We first explain how LLMs encode information in their parameters and why this architecture inherently limits factual recall. We also review strategies proposed to mitigate catastrophic forgetting, including **parameter-efficient fine-tuning (PEFT)** and **context engineering**, and explain why they remain insufficient for exhaustive, provenance-linked queries. We then describe retrieval-augmented generation (RAG) as a partial remedy and its remaining shortcomings for exhaustive queries. Next, we introduce KGs and graph retrieval-augmented generation (GraphRAG) as a path toward complete, grounded, and deterministic scientific question answering. Finally, we survey existing plant KGs resources, outline evaluation criteria, discuss practical limitations, and propose a community roadmap for building open, continuously updated plant KGs coupled to LLM interfaces.

### How LLMs “store” information (and why weights are not enough)

LLMs are trained to predict the next chunk of text (**token**). During training, they do not build a table of facts like a database (gene IDs → functions → citations). Instead, they adjust billions of internal numbers (**weights**) so that, given a prompt, they generate text that statistically resembles what they have seen before. This makes LLM recall behave like pattern completion: they can often state common relationships, but coverage is uneven across relation types and prompts, and they cannot guarantee **complete, queryable enumerations**. Because **pre-training** optimizes fluent text prediction, rather than evidence tracking, **provenance** is not naturally encoded nor guaranteed. Finally, importantly for “keeping the model up to date,” adding new training data (e.g., newest scientific articles) can degrade or overwrite previously learned information, leading to **catastrophic forgetting** (Huang *et al*., 2024); Figure 1A. Even targeted “knowledge edits” can cause changes to other facts or broader abilities by perturbing weights, often in an unexpected and unpredictable manner (Gu *et al*., 2024)(Figure 1B). Consequently, this results in **parametric knowledge** that **drifts** with training data and is **inherently incomplete,** often making efforts to improve the model performance a Sisyphean task. They perform poorly at **exhaustive enumeration** and cannot guarantee provenance.

**Figure 1.**
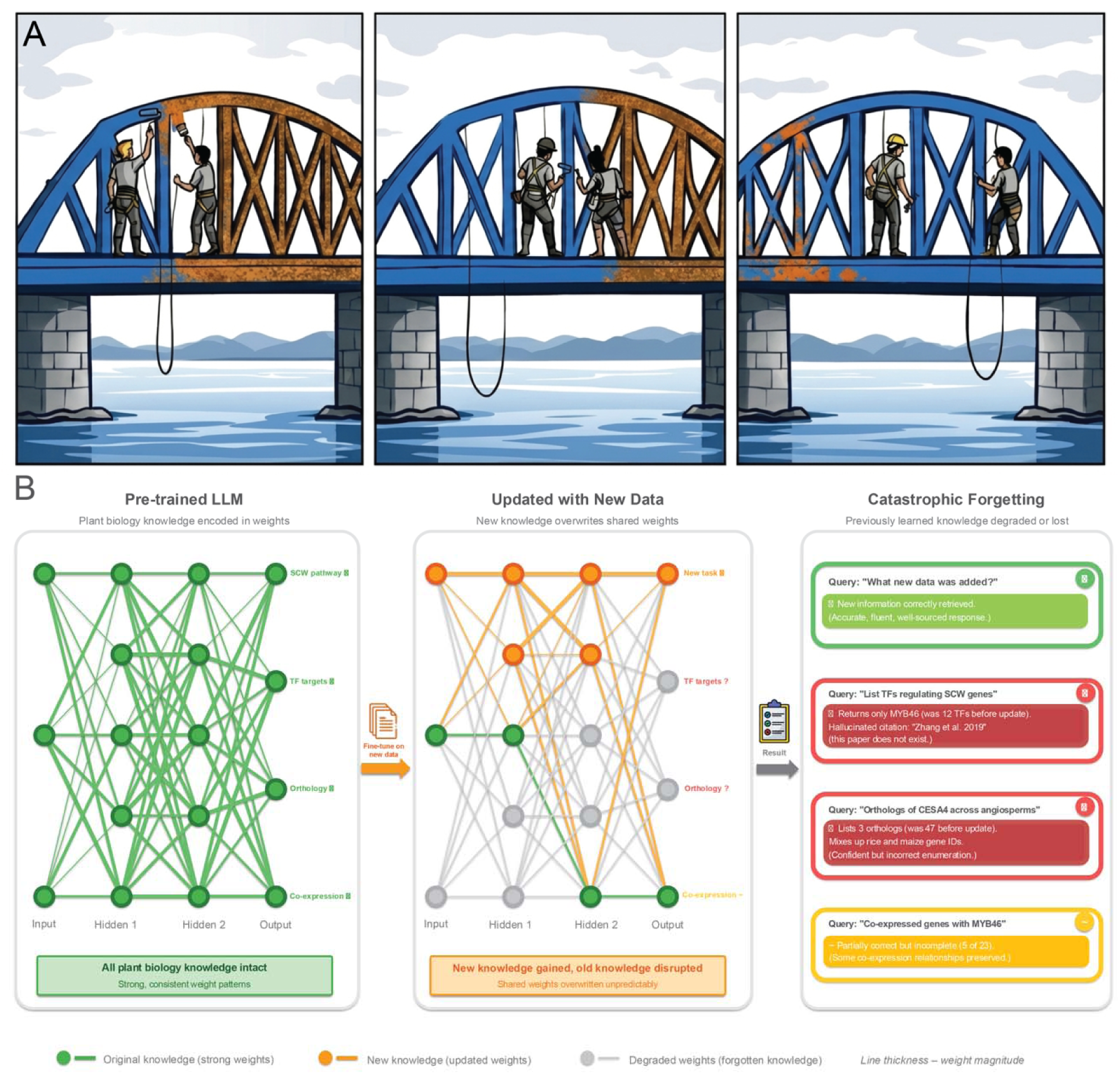
Catastrophic forgetting in large language models. A. Illustration of catastrophic forgetting. Pre-trained LLM (left panel) is updated with new information (middle panel), resulting in a loss of previous knowledge (right panel). Blue color represents paint (knowledge), while brown color represents rust (lack of information). B. A pre-trained LLM encodes plant biology knowledge (e.g., SCW pathways, transcription factor–target relationships, orthology, co-expression) as distributed weight patterns across layers (green nodes and connections). Line thickness is proportional to weight magnitude; all four output domains are reliably recalled. Fine-tuning on new data updates weights in the upper layers (orange), but because parameters are shared across tasks, previously learned weight patterns are disrupted (gray, dashed connections). The perturbation is unpredictable: some knowledge domains are severely affected while others are partially preserved. Downstream consequences for the end user: queries about newly added information succeed (top, yellow), but previously reliable queries now return incomplete enumerations (1 of 12 transcription factors; 3 of 47 orthologs), hallucinated citations, and confused gene identifiers (red). Some knowledge is partially retained but degraded (bottom, amber). These failure modes are difficult to detect because the model responds with the same confident tone regardless of answer quality.

Several approaches have attempted to reduce catastrophic forgetting during **fine-tuning**. In multi-task instruction tuning, models train on a broad mixture of tasks framed as natural-language instructions, which helps preserve general-purpose capabilities as new ones are added (Wei *et al*., 2021; Ouyang *et al*., 2022; Chung *et al*., 2024). **PEFT** methods limit forgetting by restricting which parameters change: selective methods freeze most weights and update only a small subset such as **bias terms** (Ben Zaken *et al*., 2022); **reparameterization** approaches like LoRA inject trainable low-rank matrices while the original weights stay frozen (Hu *et al*., 2021), with **quantized variants** such as QLoRA further cutting memory costs (Dettmers *et al*., 2023) and additive strategies insert **small trainable modules**, whether **adapter layers** (Houlsby *et al*., 2019) or **learnable soft prompts** (Lester *et al*., 2021; Li and Liang, 2021), without altering the base model. (Biderman *et al*., 2024) confirmed that LoRA ’learns less and forgets less’ compared with full fine-tuning, a trade-off between **adaptation depth** and **knowledge retention**. More recently, **agentic context engineering** avoids weight modification entirely by **evolving the input context of a frozen model** through **structured reflection and incremental updates** (Zhang *et al*., 2025). These strategies reduce forgetting, but none of them guarantee the exhaustive, provenance-linked enumeration that plant biology queries demand.

To demonstrate how poorly current LLMs retrieve exhaustive, citable biological knowledge, we have queried the three most popular LLMs: ChatGPT, Claude and Gemini with two questions: ‘list all of the Arabidopsis Gene Identifier (AGI) codes of transcription factors controlling SCW formation in Arabidopsis’ (testing for provenance and completion). We used a review by (Zhang *et al*., 2018) as ground truth (39 SCW transcription factors) (Figure 2A) and The Arabidopsis Information Resource (TAIR) gene alias file for AGI alias names. To note, this paper contains mainly the core factors in the SCW regulatory network, with the first layer centred around NAC transcription factors (i.e., VND1-7 and NST1-3), the second layer centred around MYB46 and MYB83 and the third layer containing transcription factors (TFs) whose expression is regulated by them. Hence, additional ground truth entries were incorporated when AGI codes generated by LLMs could be validated through supporting literature, resulting in a total of 47 SCW transcription factors.

**Figure 2.**
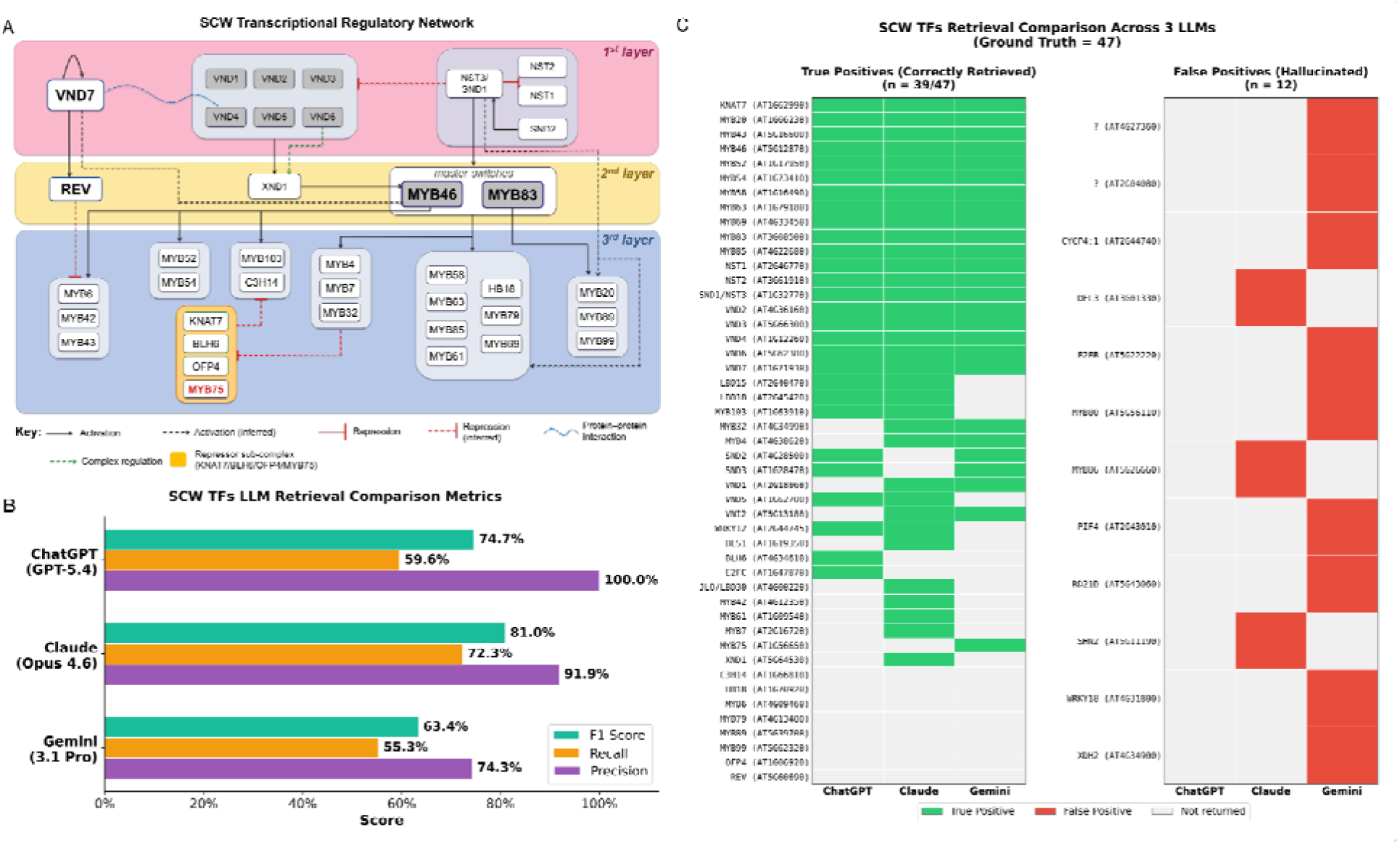
LLM recall of the SCW transcriptional regulatory network in *Arabidopsis thaliana*. (A) Curated three-tier regulatory hierarchy controlling SCW biosynthesis, compiled from published experimental evidence. Bold labels denote well-characterised master or key regulatory nodes. First-layer regulators (pink) include NAC-domain master switches (VND1–7, NST1/2, SND1/2). Second-layer regulators (yellow) include the MYB46/MYB83 master switches, REV, and the repressor XND1. Third-layer regulators (blue) comprise downstream transcription factors, including the KNAT7/BLH6/OFP4/MYB75 repressor sub-complex (orange box). MYB75 is shown in red to denote its repressive activity (Zhang *et al*., 2018). Edge types indicate experimentally validated activation (solid arrows), inferred activation (dashed arrows), experimentally validated repression (solid red flat-headed lines), inferred repression (dashed red flat-headed lines), protein–protein interactions (blue dotted curves), and complex-mediated regulation (green dashed arrows). (B) Barplot of F1 score, recall and precision for three large language models (ChatGPT: GPT-5.4, Claude: Opus 4.6, and Gemini: Gemini 3.1 Pro) tasked with retrieving AGI codes of transcription factors involved in SCW formation in Arabidopsis thaliana. (C) Responses from three LLMs (ChatGPT, Claude, Gemini) to the query *"List all AGI codes of transcription factors controlling SCW formation in Arabidopsis."* AGI codes matching the ground truth of 47 SCW transcription factors are shown in green (true positives), while AGI codes not present in the ground truth are shown in red (false positives). ‘?’ was assigned if AGI code is not in The Arabidopsis Information Resource (TAIR) gene alias file.

We used API calling to generate the answers based on the latest models GPT-5.4 (OpenAI), Gemini-3.1-preview (Google) and Opus-4.6 (Anthropic). After expanding the ground truth to 47 genes based on literature verification, Claude achieved the highest F1 score (81.0%), driven by the best balance of recall (72.3%) and precision (91.9%) among the three models. ChatGPT attained perfect precision (100.0%) but at the cost of lower recall (59.6%), yielding an F1 score of 74.7%. Gemini scored lowest across all three metrics (F1 = 63.4%, recall = 55.3%, precision = 74.3%), returning the most false positives and fewest true positives (Figure 2B). These results suggest that Claude offers the most reliable trade-off between retrieving known SCW transcription factors and avoiding spurious entries, while ChatGPT’s conservative output minimises false positives but misses a substantial portion of the network.

Among the false positives, several patterns of LLM failure emerge. First, some genes returned by Gemini are not transcription factors at all: AT2G04080 encodes a MATE efflux transporter, CYCP4;1 (AT2G44740) is a cell cycle cyclin, XDH2 (AT4G34900) is a metabolic enzyme and RD21B (AT5G43060) is a cysteine protease. Similarly, Gemini returned WRKY18 (AT4G31800), a defence-related TF, likely confusing it with the SCW repressor WRKY12 (AT2G44745). These patterns, in which the model returns non-TF proteins and confusing gene name aliases, illustrate the fundamental limitation of LLMs for biological fact retrieval: they learn statistical associations between words and concepts during training rather than querying structured databases. Next, the false negatives show 6/8 genes missed by all three models present in the 3rd layer (i.e. MYB6, C3H14, HB18, MYB79, MYB89 and MYB99), while REV (AT5G60690) and OFP4 (AT1G06920) are found in the 2nd layer and repressor sub-complex, respectively. This means that all three models reliably recall the well-characterised 1st-layer NAC master switches (VND1–7, NST1/2, SND1) and the prominent 2nd-layer MYB regulators (MYB46, MYB83), but systematically fail to retrieve genes that sit further downstream in the regulatory hierarchy. Notably, among the missed 3rd-layer genes, C3H14 and HB18 belong to non-MYB transcription factor families, suggesting that LLMs are biased towards the frequent mentions of “NAC–MYB" in SCW review literature. The remaining four (MYB6, MYB79, MYB89, MYB99) are less frequently cited MYBs compared to their well-retrieved counterparts such as MYB58, MYB63 and MYB85. Together, these results indicate that LLM recall scales with literature frequency, regardless of their validated roles in the SCW network.

Thus, because of the resource-intensive pretraining, the fact that existing fine-tuning strategies cannot guarantee completeness or provenance, and the issues with retrieving exhaustive, factual knowledge, as illustrated by the poor recall observed across all three models (Figure 2B), standalone LLMs are unsuitable for queries that require complete enumeration with provenance, a common need in plant biology. Addressing this limitation requires decoupling the language and reasoning capabilities of LLMs from the knowledge they access.

### Retrieval-augmented generation as a bridge to verifiable LLMs

**Retrieval-augmented generation (RAG)** is a pragmatic way to address the lack of provenance in standalone LLMs. The core idea is to separate language ability and reasoning from knowledge storage: instead of relying on storing the knowledge in weights, a RAG system first retrieves relevant evidence from an external **corpus** (e.g., papers, manuals, curated documents), then asks the LLM to answer using only that evidence. Earliest attempts combine parametric memory (the LLM) with non-parametric memory (a searchable document index), improving performance on knowledge-intensive tasks and enabling updates by changing the document index rather than retraining the model (Lewis *et al*., 2020).

A typical RAG pipeline has four conceptual steps. First, you define the “allowed knowledge” by curating a **corpus** (for example, a set of review articles, lab protocols, or a frozen snapshot of a database). Second, documents are split into smaller passages (“chunks”) and converted into numerical representations (“**embeddings**”) stored in a **vector index**. Third, when the user asks a question, a **retriever** converts the question into an embedding vector and searches the index to return the top-k most relevant passages. Fourth, the LLM generates an answer conditioned on those passages, ideally attaching citations to the retrieved sources. This simple pattern matters scientifically because it provides three benefits that a closed-book LLM cannot guarantee: (i) **completeness** (i.e., by incorporating new documents), (ii) **provenance** (i.e., by showing which text supports the answer , and (iii) reduced pressure to “guess” when the model does not know, resulting in fewer hallucinations (Lewis *et al*., 2020).

RAG improves **provenance**, but it still does not guarantee exhaustive enumeration for "list everything" questions. In most implementations, retrieval returns only a small top-k set of passages (e.g., 5–50) that are most similar to the query, and the system generates an answer from that limited evidence window. Even if the underlying corpus contains hundreds or thousands of relevant items scattered across many papers, retrieving and synthesizing all of them is infeasible for two reasons. First, retrieval performance often saturates or degrades as more passages are added, because the LLM struggles to attend to relevant information faithfully over very long concatenated contexts (Izacard and Grave, 2021), due to their limited context window size. Second, the computational cost is prohibitive: LLMs consume resources for every input token read, every output token generated, and in modern reasoning models, every reasoning token. While per-token prices have fallen dramatically and model capabilities have increased, processing hundreds of full-text papers to answer a single user query remains economically impractical. The most pragmatic path, therefore, is to extract structured facts from the literature once, store them with provenance, and reuse them for many downstream queries, which we discuss next.

### Crystallized information with knowledge graphs provides a path to complete, grounded and exhaustive chatbots

KGs offer a complementary strategy: instead of repeatedly re-reading thousands of documents at query time, we crystallise facts into a structured store that can be **queried deterministically**. A KG represents knowledge as a network of nodes (entities) and edges (relations, Figure 3). In the simplest view, each edge is a statement of the form (subject, predicate, object), for example: (MYB46, regulates, CESA4) or (AT1G01010, has_GO_term, “SCW biogenesis”). The key difference from free text is that these statements can carry identifiers (gene IDs, ontology IDs), **typed relations**, and metadata, which most importantly contains provenance (which paper, which database, which sentence and which experiment). This “**executable structure**” makes KGs behave like a database you can easily query, rather than a collection of papers that you have to read.

**Figure 3.**
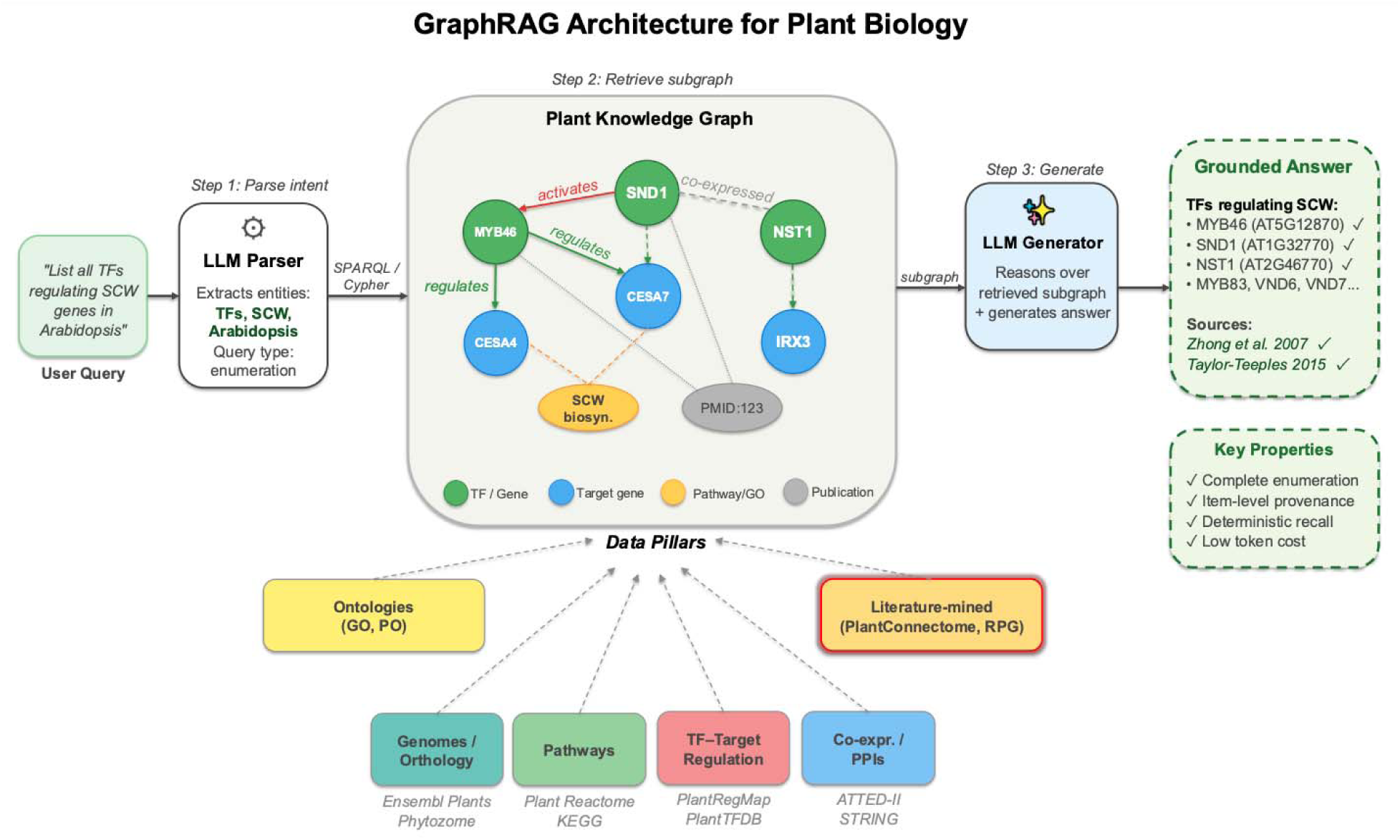
Graph retrieval-augmented generation architecture (GraphRAG) for plant biology. Top: the query pipeline proceeds in three steps. (Step 1) An LLM parser extracts entities and intent from the user’s natural-language query (e.g., transcription factors, SCW, *Arabidopsis*; query type: enumeration). (Step 2) The parsed entities are used to retrieve a relevant subgraph from a plant KG via structured query languages (SPARQL, Cypher). The KG (center) stores biological entities as typed nodes: genes and transcription factors (green), target genes (blue), pathway and Gene Ontology terms (yellow), and publications providing provenance (gray), connected by explicitly labeled relations (e.g., *regulates*, *activates*, *co-expressed*). (Step 3) The LLM generator reasons over the retrieved subgraph and produces a grounded answer containing complete gene lists with identifiers and item-level citations. Bottom left: the six data pillars feeding the KG, spanning curated databases (Ensembl Plants, Phytozome, Plant Reactome, KEGG, PlantRegMap, PlantTFDB, ATTED-II, STRING), ontologies (GO (Gene Ontology), PO (Plant Ontology)), and literature-mined resources (PlantConnectome, RPG). The literature-mined pillar is highlighted (red border) as it is the primary source of regulatory and interaction data for the knowledge graph. Center right: Key properties of the GraphRAG approach are listed: complete enumeration within the defined scope, item-level provenance linking each returned entity to its supporting database record or publication, deterministic recall (identical queries return identical results), and low token cost (the LLM processes a compact subgraph rather than re-reading source documents).

**GraphRAG** is a system in which the “memory” you retrieve is a graph of entities and relations, and the LLM answers using only that graph (Edge *et al*., 2025, Preprint). When a user poses a question, the system uses the KG to retrieve a relevant subgraph (e.g., all transcription factors and target genes involved in a given pathway, along with their attributes and source references). The LLM can then reason (e.g., which part of the retrieved information is relevant) over this subgraph and generate a final answer in fluent English, grounded in the retrieved facts. Because the answer is based on authoritative database content rather than the LLM’s parametric memory, and the number of tokens needed to process the retrieved graph is much less than re-reading the corresponding papers, GraphRAG can be both complete and trustworthy. Recent studies in specialized domains support this approach: incorporating domain KGs into the Question and Answer pipeline has been shown to improve answer accuracy and coverage while suppressing hallucinated content (Deng *et al*., 2025; Wu *et al*., 2025b). For example, a Crop GraphRAG system (integrating a pest and disease KG for crops) produced more precise answers and returned more of the relevant facts compared to an LLM alone, and it “effectively suppresses hallucinated content” in a specialized agricultural context (Wu *et al*., 2026). These successes illustrate the potential of GraphRAG for any knowledge-intensive field.

### Knowledge graphs make it easy to integrate different data types

A biological KG can be built by integrating multiple heterogeneous data sources into a shared identifier space and a common set of typed relations. In practice, curators (or pipelines) ingest structured databases (e.g., pathways, gene/protein annotations, chemicals, ontologies), interaction resources (protein-protein interactions (PPIs), genetic interactions), regulatory resources (TF–target edges, cis-elements), and experimental summaries (expression, phenotypes), then normalize entities to stable IDs (e.g., UniProt/Ensembl/NCBI, GO/PO, ChEBI (Chemical Entities of Biological Interest)) and represent assertions as triples (subject–predicate–object) with provenance. This “linked data” approach is exemplified by Bio2RDF (Belleau *et al*., 2008), which converts dozens of public bioinformatics databases into interoperable Resource Description Framework (RDF) with normalized URIs (Uniform Resource Identifiers) and shared semantics, enabling cross-dataset querying. Likewise, heterogeneous biomedical networks such as Hetionet (Himmelstein *et al*., 2017) explicitly combine many node/edge types across domains to support integrative inference and querying. In plant science, Plant Reactome (Naithani *et al*., 2020) illustrates how curated pathway knowledge can be connected to genomes and orthology-based projections across many species, providing an integrated backbone that can be linked onward to other resources. Finally, literature seamlessly complements these data sources: systems like INDRA (Gyori *et al*., 2017) assemble mechanistic “statements” by combining text-mined assertions with structured pathway/database knowledge into standardized causal relations, again with evidence tracking.

### Examples of knowledge graphs in plant sciences

In plant biology, many such data resources exist, though they are often siloed. For example, AgroLD is a large KG that integrates over 1.08 billion factual triples from ∼151 datasets covering plant genomes, proteins, pathways, gene regulation datasets (e.g. PlantTFDB, PlantRegMap), protein–protein interactions, and more (Pierre *et al*., 2025). Beyond integrative, ontology-driven KGs, a growing class of LLM-coupled plant KGs is emerging, where models either (i) extract structured statements from literature into a graph, and/or (ii) perform graph-grounded QA (Question Answering). For example, PlantScience.ai builds and maintains the Plant Science Knowledge Graph (PSKG) via an automated LLM-driven extraction pipeline (AutoSKG) to enable source-traceable querying over plant concepts and relations (Yu *et al*., 2026). In crops, SeedLLM·Rice integrates a domain LLM with a Rice Biological Knowledge Graph (RBKG) that consolidates curated genome annotations and related knowledge to improve reasoning and retrieval for rice genetics and breeding questions (Yang *et al*., 2025). In photosynthesis, LLM-based pipelines have been used to construct and evaluate a dedicated literature-derived KG (validated on ∼5,000 photosynthesis articles) and to support KG-structured responses in specialized assistants(Yoon *et al*., 2025, Preprint). Together with large-scale extraction KGs such as PlantConnectome (LLM-mined relations from >71,000 plant articles) (Lim *et al*., 2025) and broad linked-data efforts like AgroLD (Pierre *et al*., 2025), these examples illustrate an accelerating shift from siloed databases toward LLM-assisted knowledge acquisition + graph-native integration and querying.

### Open challenges and a roadmap towards GraphRAG-powered chatbots

Realizing GraphRAG for plant biology requires solving several interlinked data and governance problems. The first is **entity disambiguation**: gene symbols, accession numbers, metabolite names, and gene family labels are context-dependent and must be normalized to stable identifiers to avoid fragmented or conflated nodes in the graph. Existing resources such as TAIR (Rhee *et al*., 2003), Phytozome (Goodstein *et al*., 2012), and Ensembl Plants (Kersey *et al*., 2018) provide stable identifiers for many model and crop species, but coverage remains uneven for non-model organisms, and text-mined entities extracted from literature frequently lack standardized identifiers altogether (Bachman *et al*., 2018). The second is **relation normalization**: different databases encode similar biological relationships using incompatible vocabularies (e.g., "regulates," "activates transcription of," and "positively regulates expression" may all describe the same edge), and mapping these onto a controlled predicate schema is essential for list-style queries to yield closed, deduplicated result sets (Han *et al*., 2025, Preprint). Third, evidence grading must be encoded at the level of individual assertions, specifying which experiment, figure, species, tissue, and condition supports each edge, while allowing contradictions to coexist rather than silently overwriting one another. This is exemplified by PlantConnectome, which shows the type of evidence underpinning an edge (Lim *et al*., 2025).

A practical roadmap towards GraphRAG-powered chatbots is therefore to (i) identify shared identifier and predicate standards, building on community frameworks such as the Bioregistry (Hoyt *et al*., 2022) and OBO (Open Biological and Biomedical Ontology) Foundry (Smith *et al*., 2007); (ii) build open, testable normalization pipelines that resolve synonyms, map legacy accessions, and flag ambiguous entities; (iii) attach machine-queryable provenance to every assertion, so users can filter results by evidence type and confidence; (iv) release versioned, citable KG snapshots with continuous update logs, making queries reproducible across time; and (v) benchmark "list all X" queries against curated gold standards (e.g., expert-validated transcription factor lists for defined regulatory networks, complete enzyme sets for specific biosynthetic pathways across species) using existing plant KGs as scaffolds. The GraphRAG interfaces built on these resources should be developed as open-source tools with transparent retrieval logic, so that users can inspect the subgraph underlying any answer and trace each returned entity to its supporting evidence. Such transparency is not merely a technical convenience but a prerequisite for scientific trust.

## Conclusion

Plant biology knowledge is unusually “graph-shaped.” We routinely reason in terms of entities (genes, proteins, metabolites, traits, tissues, stresses) and relationships between them (orthology, regulation, pathway membership, physical interaction, localization, phenotypes). This makes the plant community well-positioned to benefit from KGs as a shared, queryable substrate for integrating literature and multi-omics resources. Coupled to LLMs via GraphRAG, such graphs can underpin scientific assistants that return exhaustive enumerations within a defined scope, with item-level provenance linking each gene/edge to its supporting database record and/or publication. This would enable routine retrieval of complex evidence spanning hundreds of papers and heterogeneous genomic/omic datasets, while preserving the audit trail required for scientific use. By crystallizing plant knowledge into provenance-linked graphs and placing LLMs on top via GraphRAG, we can turn “read 1000 papers” into a deterministic query, thereby delivering exhaustive, evidence-backed answers at a reasonable query duration.

#### Box 1. Definitions

| Key | Definition |
| --- | --- |
| Agentic context engineering | A framework that treats contexts as evolving playbooks that gradually accumulate new strategies, refine existing ones, and keep them well organized through a modular cycle of generation, reflection, and curation. |
| Context adaptation | A method to adapt a frozen LLM by iteratively refining the input prompts or context using structured feedback and updates. No model weights are changed; improvements come entirely from manipulating the input instructions, strategies or evidence. |
| Adaptation depth | The number of layers or parameters in a model that are modified during fine-tuning. Greater adaptation depth affects more of the model's internal representations but increases risk of forgetting previously learned knowledge. |
| Adapter layers | Small, lightweight components added into a pre-trained language model between its existing internal layers. When adapting the model to a new task (e.g., extracting plant-gene relationships), only these small components are trained while the rest of the model stays unchanged. |
| Bias terms | Trainable parameters added to the weighted sum of neuron inputs before activation, controlling the baseline output of neurons. Updating only bias terms is a lightweight way to adapt a model without changing its core weights. |
| Catastrophic forgetting | When learning new data/tasks causes performance or knowledge on earlier data/tasks to degrade. |
| Complete, queryable enumerations | E.g., all enzymes in a pathway across taxa, or all TF→target edges in a defined scope |
| Completeness | The ability to retrieve an exhaustive and correct set of all relevant entities for a given query. |
| Corpus | The collection of documents you allow the system to use as |
|  | evidence. |
| Drifts | Changes in stored knowledge as model parameters are updated during training or fine-tuning. |
| Embeddings | A numeric representation of text used to compute similarity. Similar words and sentences would produce numerically similar, correlated vectors. This allows rapid identification of semantically similar words, sentences, and documents by computing, e.g., the cosine similarity between their vectors. |
| Entity disambiguation | The process of resolving ambiguous names to unique, stable identifiers, so that entities mentioned in different sources or contexts are correctly recognized as the same or distinct entries in the knowledge graph. For example, "Arabidopsis thaliana," "thale cress," "Arabidopsis," and NCBI taxonomy ID 3702 must all map to the same organism node. |
| Evolving the input context of a frozen model | Iteratively modifying prompts or preceding input text to improve a frozen model's outputs, rather than changing the model weights. |
| Executable structure | Executable structure (in the context of knowledge graphs) means the knowledge is stored in a machine-interpretable form, so a computer can run formal queries and reasoning over it, rather than "reading" prose and guessing intent. |
| Exhaustive enumeration | A systematic, complete retrieval of all entities that satisfy a defined query within a given scope — for example, all transcription factors regulating a specific pathway in a specific organism — such that no known qualifying entity is omitted from the result set. |
| Fine tuning | A process in which a subset or all of the weights of a pre-trained language model, which has already learned general knowledge from large amounts of text, is further trained on a smaller, domain-specific dataset, adapting the model's learned representations to a particular task. |
| GraphRAG | LLM + graph-based retrieval + evidence-linked generation, designed to return complete, grounded, queryable answers, especially for list-style, multi-hop biological questions. |
| Inherently incomplete | Lacking guarantees of full coverage due to reliance on probabilistic pattern learning rather than structured data storage. |
| Knowledge retention | The ability of a model to maintain previously learned information after fine-tuning or adaptation. High retention means new learning does not overwrite existing knowledge. |
| Learnable soft prompts | Trainable input vectors used to guide the outputs of a frozen model. Unlike hard-coded prompts, they are optimized during training to improve task performance without changing model weights. |
| Parameter-efficient fine-tuning (PEFT) | A set of strategies to adapt a model to new tasks by modifying only a small subset of its parameters, rather than performing full fine-tuning on all weights. This reduces memory use and limits catastrophic forgetting. |
| Parametric knowledge | Information implicitly encoded in model parameters during training, rather than stored in an explicit, queryable database. |
| Pre-training | Large-scale training on broad text to learn general language/statistical patterns.<br>In LLMs, this is the most resource-intensive and expensive step, often requiring millions of dollars. |
| Provenance | The evidence trail for a claim (paper/database record supporting a statement or edge). |
| Quantized variants | Model versions in which weights are stored in lower numerical precision (e.g., 8-bit instead of 32-bit) to reduce memory and computational requirements, often combined with PEFT methods like LoRA. |
| Queried deterministically | Given the same data, the same query, and the same query rules, the system will return the same result set every time by applying explicit logic rather than probabilistic generation. |
|  | LLMs are probabilistic text generators, and tend to give different answers even when the same query is used. |
| Relation normalization | The process of mapping diverse linguistic and database-specific descriptions of biological relationships onto a controlled, standardized set of edge types, so that semantically equivalent assertions from different sources can be queried together. |
| Reparameterization | A strategy that re-expresses a model's existing parameters in an alternative, often lower-dimensional form, so that fine-tuning can be performed in a more compact space. This reduces the number of trainable parameters while preserving the model's capacity. The adapted parameters can typically be merged back into the original model with no additional cost at inference. |
| Retrieval-Augmented Generation (RAG) | A system that retrieves external evidence and conditions LLM generation on it. |
| Retriever | Component that selects candidate passages given a query. |
| Small trainable modules | Compact components inserted into a model, such as adapter layers or soft prompts, that can be trained while leaving the base model unchanged. |
| Structured reflection and incremental updates | A process where outputs are analyzed and fed back to refine the model's input context or prompts step by step. Structured reflection provides organized feedback, and incremental updates gradually improve behavior without modifying weights. |
| Token | A unit of text that the model processes (often part of a word). |
| Top-k | k is the number of retrieved passages provided to the generator. |
| Typed relations | Typed relations are edges with explicitly defined predicate labels (e.g., regulates, ortholog_of, part_of), enabling unambiguous interpretation and precise querying of biological relationships. |
| Vector index | Database that supports similarity search over embeddings. |
| Weights | Or parameters: The learned internal numbers of the neural network. |

